# Understanding malaria transmission in Angola: first detection of *Plasmodium malariae* in *Anopheles spp*. and the role of secondary vectors in the Balombo region, Benguela Province

**DOI:** 10.64898/2026.09.14.751460

**Authors:** Quentin Narpon, Jean-Claude Toto, Estelle Lucarz, Maria Adelaide Dos Santos, Almeida Ingles, Clarisse Girault, Patrick Besnard, António Manuel Cabinda, José Franco Martins, Anne Poinsignon, Sylvie Manguin

## Abstract

**Background:** Angola is one of the most affected countries in sub-Saharan Africa and was off-track to meet the WHO Global Technical Strategy targeting of a 75% reduction in malaria incidence and mortality by 2025. Recent studies highlighted major knowledge gaps regarding malaria vectors in Angola, particularly secondary vectors. A malaria control project initiated in 2007 in Balombo (Benguela Province, central Angola) has monitored malaria dynamics for 18 years, revealing increased *Plasmodium* prevalence and more frequent detection of non-falciparum species, mainly *Plasmodium malariae* (in co-infection with *P. falciparum*) in villagers of the Balombo region. This study aimed to characterize *Anopheles* populations in this region by assessing species composition, behavioural patterns, and infection rate. And to a less extent, to compare the performance of two indoor and outdoor mosquito trapping methods.

**Methods:** Surveys were conducted during the rainy (December 2023 and 2024) and dry seasons (July 2024) in villages around the town of Balombo. Adult mosquitoes were collected indoors and outdoors using CDC Light Traps and BG-Pro traps in randomly selected houses. Female *Anopheles* were morphologically and molecularly identified. Primary and secondary vectors were screened for *Plasmodium spp*. using qPCR-high resolution melting, and positive samples were sequenced.

**Results:** A strong seasonal effect was observed in *Anopheles* populations, with higher abundance, diversity, and *Plasmodium*- infection rates during the rainy season. *An. funestus* remained the main vector, but *Plasmodium spp.* was also detected in five other species: *An. arabiensis*, *An. gambiae*, *An. marshallii*, *An. rufipes*, and *An. maculipalpis*. This is the first report of *P. malariae* in *Anopheles* mosquitoes and the first mention of the role of secondary vectors in malaria transmission in Benguela Province.

**Conclusion:** This study provides a thorough and updated inventory of *Anopheles* species in the Balombo region. Molecular biology confirmed the presence of *P. falciparum* and *P. malariae* in several species of *Anopheles* underscoring the existence of a complex transmission system in villages of Balombo. Infected exophagic and exophilic *Anopheles* species, exhibiting opportunistic behaviour, likely contribute, as secondary vectors, to the increase of *Plasmodium spp.* transmission in the study sites. Therefore, confirming the need to implement complementary interventions targeting outdoor malaria transmission.

**Author Summary:** This study provides a thorough and updated inventory of *Anopheles* species in the Balombo region, conducted nearly two decades after the implementation of vector control interventions, in a setting where LLINs continue to be distributed. Sequencing confirmed the presence of *Plasmodium spp.* in *Anopheles* during both the dry and rainy seasons and revealed the first detection of *P. malariae* in mosquitoes collected in Angola. Although *An. funestus* remains the primary malaria vector in the Balombo region, the detection of *Plasmodium spp.* in five other *Anopheles* species highlights the existence of a more complex transmission system across the studied villages. While the sampling protocol and the relatively low numbers of *An. funestus* captured in some villages preclude definite conclusions regarding its behaviour, the possibility of a shift towards exophagy and exophily in some villages of Balombo is possible. It is therefore necessary to step up regular entomological surveillance, particularly in villages that were the focus of vector control campaigns between 2007 and 2011. Moreover, the detection of *Plasmodium spp.* in several secondary vectors exhibiting opportunistic and exophilic behaviours, such as *An. marshallii*, suggests that LLIN coverage alone may be insufficient to prevent exposure to infectious mosquito bites. A comprehensive assessment of LLINs use throughout the year is therefore needed, together with the adaptation of vector control strategies to address the contribution of exophilic vectors to malaria transmission in the villages of Balombo.

## Introduction

Despite major global advances in the fight against malaria between 2000 and 2015, which led to a significant decline in the number of cases and deaths, the following decade was marked by stagnation, or even an increase, in these indicators in several regions of the world. In 2024, sub-Saharan Africa accounted for 94% of both malaria cases (282 million), and deaths (estimated at 610,000) worldwide [1]. Angola is one of the most malaria-affected countries in the region, with nearly 10 million cases and 16,000 deaths recorded in 2024, placing it 6^th^ and 8^th^ globally for malaria cases and deaths, respectively [1]. The country was off-track to meet the 2025 milestone of a 75% reduction in malaria incidence and mortality set by the World Health Organization (WHO) in its Global Technical Strategy for malaria 2016-2030 [1]. Indeed, no significant progress has been observed in reducing either malaria incidence or mortality compared with 2015 levels. On the contrary, the malaria incidence even increased by 50% between 2015 and 2024, placing Angola second (after Sao Tome and Principe) among the Central African countries with the highest increase in incidence over the past decade [1]. Nonetheless, it should be noted that Angola made significant progress in reducing its malaria mortality rate, which declined by half, from 58.2% in 2016 to 26.3% in 2019 [2]. However, disruption of the Angolan public health system following the COVID-19 pandemic contributed to the increasing number of deaths observed between 2020 and 2022, with an estimated number rising from 15,251 in 2019 to 20,493 and 20,286 in 2021 and 2022, respectively, before declining to 16,169 and 16,385 deaths in 2023 and 2024, respectively [3].

The exceptional stable endemicity of malaria in sub-Saharan Africa is partly due to a vectorial system, which produces inoculation rates far higher than the minimum necessary to saturate human populations [4]. It is well known that sub-Saharan Africa has the most effective and efficient vectors of human malaria parasites [5]. The four *Anopheles* species considered as the primary malaria vectors, in alphabetical order, are *Anopheles arabiensis*, *An. coluzzii*, *An. funestus,* and *An. gambiae*, due to their high levels of anthropophagy, endophagy, and endophily [5–7]. Other major vector species are also often cited in the literature, such as *An. melas* and *An. merus* (members of the *An. gambiae* complex), distributed along the western and eastern African coasts respectively, as well as other species like *An. moucheti* and *An. nili* [5].

The decline in malaria cases and deaths until up 2015 has been largely attributed to the success of vector control strategies based on the large-scale distribution of long-lasting insecticidal nets (LLINs) and indoor residual spraying (IRS) [8] that are very effective to target indoor biting and resting *Anopheles* vectors, and thus reduce malaria transmission from primary vectors [9]. Behavioral plasticity of certain vector species, such as *An. arabiensis*, which exhibits strong tendency toward zoophily and exophagy, is well documented and is thought to reduce the effectiveness of indoor-based control interventions [10]. However, the limitations of these control strategies are mostly attributed to the emergence of insecticide resistance (IR) and shifts in the behaviour of primary vector populations [9,11]. The widespread and extensive use of pyrethroids in agriculture[12], as well as their presence in most LLINs has contributed to the emergence of resistance through a range of mechanisms, including mutations at target sites, cuticular and metabolic resistances [13]. While the use of LLINs could have selected populations of primary vectors exhibiting zoophagic, exophagic, and exophilic tendencies [14,15], it has also been described that control measures could shape local *Anopheles* species composition by favouring those whose ecological and behavioural traits reduce their contact to indoor interventions [16]. The term of secondary vectors is used to describe *Anopheles* species that are known (or suspected) to play a minor or seasonal role in malaria transmission, accounting for <5% of the total malaria transmission in Africa [17]. As specified by numerous authors, their role in malaria transmission is often time and site specific [17,18]. Fewer than 15 species are consistently reported as secondary malaria vectors in sub-Saharan Africa, with *An. coustani*, *An. pharoensis*, *An. marshallii*, *An. rufipes*, *An. squamosus* and *An. ziemanni* among the most frequently cited [7,17,19]. However, the number of *Anopheles* species reported as naturally infected with *Plasmodium* spp. continues to expand [20]. Most non-dominant local vectors exhibit exophilic, exophagic, and zoophilic behaviours, enabling them to avoid contact with LLINs and thus potentially sustain malaria transmission [9,17].

Recent studies have sought to identify the challenges Angola is facing toward malaria control, with the aim of proposing more appropriate solutions [21,22]. Multiple factors related to socioeconomic conditions and healthcare system weaknesses have been implicated, including inconsistent and inadequate financial investments or limited access to healthcare in rural and remote areas [21]. Among factors directly related to vectors, the increasing prevalence of insecticide resistance, the improper use of LLINs, and the lack of resources allocated to vector control, have also been cited [22]. However, little to no mention is made about behavioural shifts in the main *Anopheles* vectors or about the diversity of species involved in malaria transmission in Angola. This lack of information is well recognized and has been underscored by several authors [21–24]. Although recent entomological studies conducted in Angola, including in Benguela Province, have focused primarily on major vectors such as *An. funestus* and members of the *An. gambiae* complex, they have consistently emphasized the need to further investigate the potential contribution of less-studied *Anopheles* species to malaria transmission in the country [23,24]. At the request of the Angolan National Malaria Control Program (NMCP), a long-term vector control program was implemented in 2007 in eight villages surrounding the town of Balombo (Benguela Province) to compare the efficacy of four indoor vector control methods. This program resulted in considerable reductions of all three indicators analysed: entomological (82.4% reduction in *Anopheles* densities), parasitological (54.8% reduction in blood smear positivity) and immunological (reduced IgG responses to *Anopheles* salivary antigens) [25]. Since then, annual parasitological surveys have been conducted by the members of Malaria Control Program (MCP) of the Sonamet Company in the same villages, and the data collected have been reported in several publications [26–28]. Results from MCP annual reports have shown a gradual increase in malaria prevalence in villagers from the Balombo region since 2016, but also a more frequent detection of non-*falciparum* infections since 2018. The first detection of *Plasmodium malariae* in the human population of Balombo, reported in the only published study on the subject to date, is based on data collected between 2007 and 2011. At that time, *P. malariae* was detected in 0.4% of samples, and in 0.8% of mixed infections with *P. falciparum* [29]. *P. malariae* is a neglected human malaria parasite, partly due to milder clinical manifestations, lower parasite densities, and frequent misidentification during routine microscopic examination, even though it accounts for nearly 10% of malaria infections in sub-Saharan Africa [30,31]. Despite a few entomological surveys occasionally conducted by the MCP and a published paper briefly describing *Anopheles* species [25], information on vectors in Balombo remains scarce, and vector species responsible for transmitting non-*falciparum* malaria have yet to be identified. Moreover, correct identification of the local malaria vectors is a key-step in defining vector control strategies [17].

The objective of this study was to characterize *Anopheles* populations present in the Balombo area, including both primary and secondary malaria vectors, by assessing species composition, behavioural patterns, and *Plasmodium* infection rates. In addition, the study aimed to evaluate the sampling performance of two mosquito trapping methods used for indoor and outdoor mosquito collections in terms of *Anopheles* abundance and species diversity.

## Methods

### Study sites

The study was conducted in eight villages located around the town of Balombo (12°21′26″ S, 14°46′16″ E), in the Benguela highlands of central Angola (Fig. 1). Balombo (∼50.000 inhabitants) lies 150 km east of the coastal city of Lobito and 420 km south of the capital, Luanda. The area is characterized by a tropical savannah with shrub prairie (Angolan miombo woodland) at an altitude of 1,200–1,300 m. The town and surrounding villages are mainly located in small river valleys bordered by hills (1,500-1,700 m) and several high peaks (2,300-2,600m). The climate is characterized by a rainy season from November to April (mean rainfall ∼140 mm in December) and a dry season from May to October (no rain in July). Temperatures range from 17–28 °C in December to 12–32 °C in July. Over the past two decades, extensive deforestation has occurred in the area due to agriculture and livestock farming, which are the main subsistence activities of local communities.

**Figure 1:**
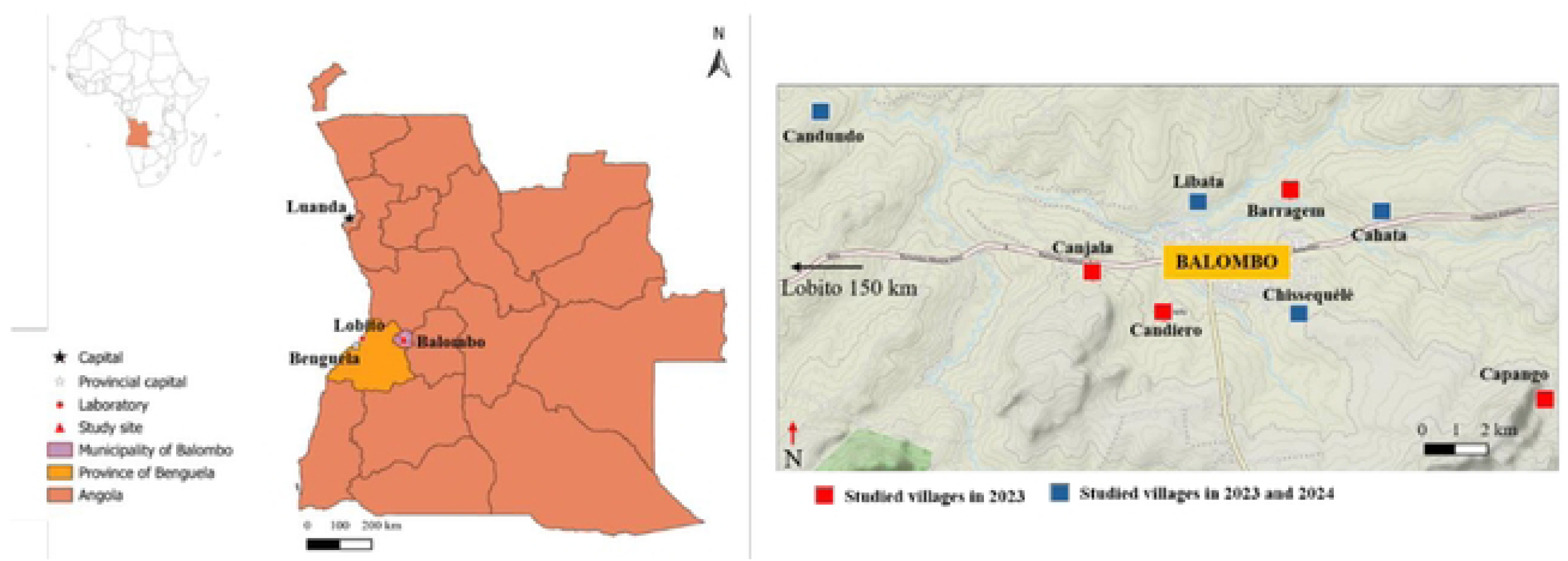
Map of Angola and mosquito collection sites in the Balombo study area.

### Adult mosquito collections

A preliminary entomological survey was conducted in December 2023 in eight villages (Fig. 1) with the objective of selecting four villages of particular interest for 2024, based on their *Anopheles* fauna and evaluating the most suitable entomological protocol for subsequent in-depth studies. Night capture was conducted in each village using several collection methods to maximize the diversity of *Anopheles* species sampled: six CDC light traps (CDC-LT) were placed indoors and outdoors in three randomly selected houses from 18:00 to 06:00 (one indoor and one outdoor per house), two Biogents BG-Pro (BGP) traps were set indoors and outdoors in a single house, and one additional BGP was installed near cattle enclosures or along passage areas for 24 h. Indoor and outdoor resting collections were also performed with a Prokopack aspirator between 05:30 and 06:00, in randomly selected houses and cattle enclosures, in each village.

Following this first survey, two consecutive nights of mosquito collection were conducted in four selected villages (Cahata, Chissequele, Candundo, and Libata) in July and December 2024 (Fig. 1, blue squares). Each village was divided into two zones; CDC-LT and BGP traps were deployed in separate zones on the first night, then swapped between zones on the second night. Traps were operated from 18:00 to 06:00, with 12 traps per night (six of each type) distributed across six randomly selected houses at least 50 m apart, with one trap placed indoors and one outdoors (the same houses were selected across both surveys). This design allowed for standardized comparison of trap performance across sites and nights, while capturing both indoor and outdoor mosquito populations. To assess potential daytime activity of *Anopheles*, five BGP traps were deployed in each of the four villages between 06:00 and 18:00 in December 2024, following the same indoor/outdoor scheme. Four BGP were installed in two of the three previously selected houses, while an additional trap was placed close to a cattle enclosure.

Collected *Anopheles* females were morphologically identified to species or complex level under a stereomicroscope using standard identification keys and the mosquito identification software developed by IRD [32]. Then, the specimens were individually preserved in tubes containing a desiccant (silica gel, Sigma-Aldrich, St. Louis, MO, USA) and stored at −20°C for molecular analyses.

### DNA extraction

Prior to DNA extraction, *Anopheles* specimens morphologically identified as primary or secondary malaria vectors were dissected, abdomen were separated from the head and thorax, and stored at −20°C. DNA from the head and thorax was extracted using the CTAB method, as described in Morlais *et al.* [33]. Briefly, samples were grounded in 200 μL of 2% CTAB solution (1 M Tris HCl pH 8.0, 0.5 M EDTA, 1.4 M NaCl, 2% cetyltrimethylammonium bromide) and homogenized with beads for 90s, and then incubated at 65°C for 10 min. Total DNA was then extracted with chloroform, precipitated with isopropanol, washed with 70% ethanol, and resuspended in DNA-free water and stored at −20°C.

### Molecular identification of Anopheles species of the An. gambiae complex and An. funestus group

Molecular identification of *An. gambiae* complex specimens was performed using the PCR–RFLP method described by Fanello *et al.* [34]. A first multiplex PCR, following the protocol in Scott et al. [35] was carried out to amplify a 390 bp fragment of the ribosomal DNA intergenic spacer region. The reaction mix contained 1 U GoTaq G2 Flexi DNA Polymerase (Promega, Madison, WI, USA), 1X GoTaq Flexi Buffer, 1.5 mM MgCl_2_, 0.2 mM of each dNTP, 0.2 µM of each primer (Table 1), and 0.5 µL of DNA template in a final reaction volume of 50 µL. The PCR was carried out with an initial step of 3 min at 94°C to activate the DNA polymerase followed by 35 cycles, each consisting of 30 s denaturation at 94°C, 30 s annealing at 56°C and 20 s extension at 72°C; the final cycle products are extended for 5 min at 72°C. After amplification, 2.5 U of Hha I, 1X Hla I Restriction Enzyme Buffer (Thermo Fisher Scientific, Waltham, MA, USA) and 4.0 µl of PCR product were directly added to the mix, and digestion was carried out at 37°C for a minimum of 3 h. Digested fragments were then run through an 2% agarose gel. Lengths of amplified species-specific products are presented in Table 1.

**Table 1.**
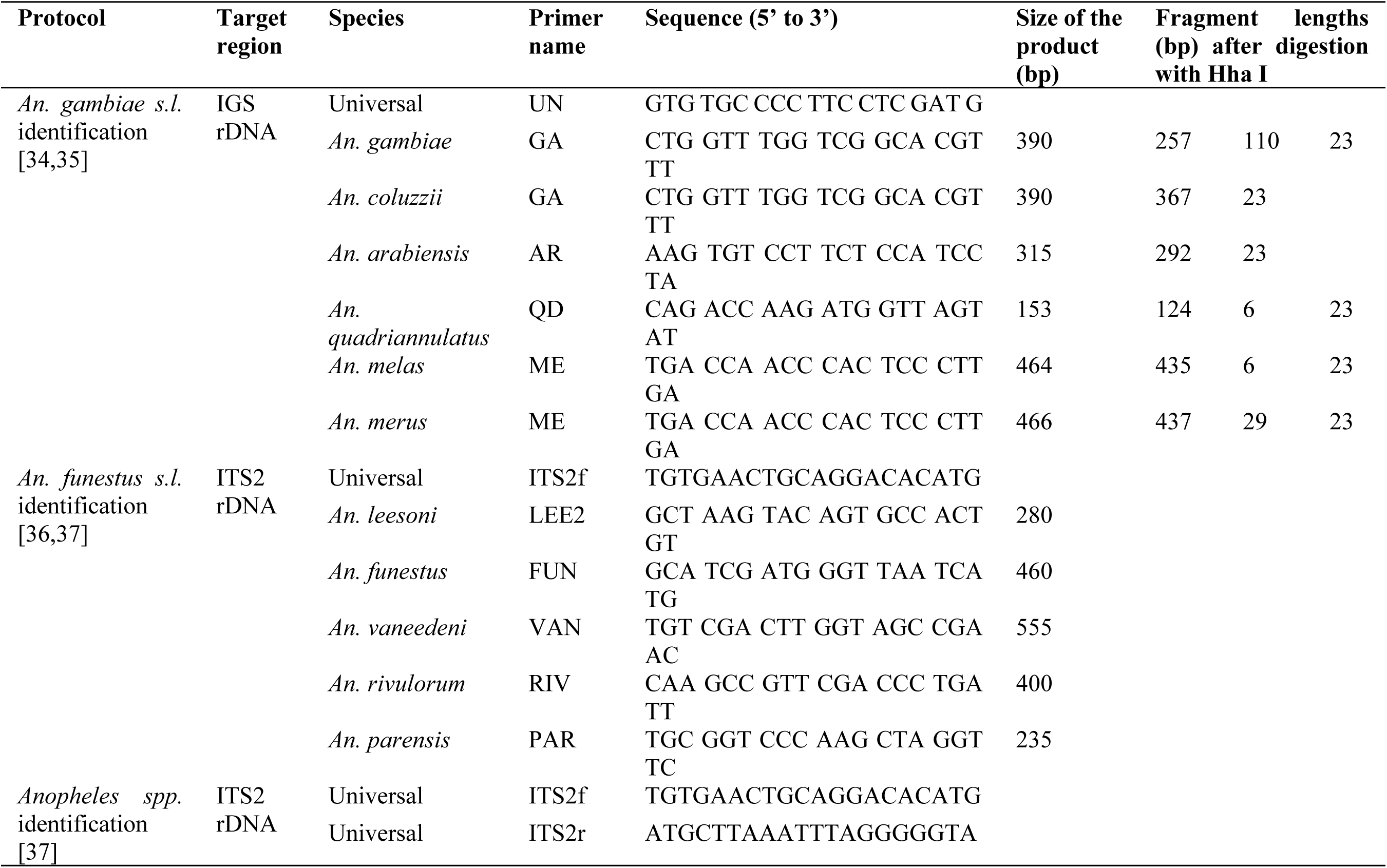
Primers used for molecular identification of *Anopheles* species, including members of the *An. gambiae* complex, the *An. funestus* group, and secondary malaria vectors.

The detection of five sub-Saharan *Anopheles* species from the *An. funestus* group was done by using a multiplex PCR described by Garros *et al.* [36], targeting Internal Transcribed Spacer 2 (ITS2) of ribosomal DNA (rDNA). PCR reactions were performed in a final volume of 25 µL containing 1 U GoTaq G2 Flexi DNA Polymerase, 1X GoTaq Flexi Buffer, 3 mM MgCl_2_, 0,2 mM of each dNTP, 0.6 µM of the universal forward ITS2 primer described by Collins and Paskewitz [37], 0.6 µM of each species-specific reverse primer (Table 1), and 3 µL of DNA template. Cycling parameters were identical to those described by Garros *et al.* [36]. The PCR products were subjected to electrophoresis on a 3% agarose gel. Lengths of amplified species-specific products are presented in Table 1.

### *Anopheles* sequencing

*Anopheles* morphologically classified as secondary vectors, *Plasmodium*-positive specimens, those with unsuccessful PCR identification, and a subset of specimens whose PCR profiles indicated less-common *An. funestus* group species, were further characterized by ITS2 rDNA sequencing. The PCR mixture contained 1U GoTaq G2 Flexi DNA Polymerase, 1X GoTaq Flexi Buffer, 1.5 mM MgCl_2_, 0.2 mM of each dNTP, and 0.2 µM of each ITS2 primers from the 5.8S and 28S coding region flanking ITS2 region (∼350-650 pb) described by Collins *and* Paskewitz [37] (Table 1). Cycling parameters were identical to those described by Garros *et al*. [36]. PCR amplification was confirmed by running PCR products on a 2% agarose gel. Size-verified ITS2 PCR products were sequenced commercially by Genewiz using Sanger method with the universal primer (forward direction). Sequence alignments were then performed within the GenBank dataset using the basic local alignment search tool BLASTN algorithm. To ensure accurate identification, BLAST results were complemented with sequence alignments in CodonCode Aligner (CodonCode Corporation, Dedham, MA, USA), particularly for *Anopheles* specimens belonging to the same species group or showing high similarity (>98–99% identity). Verified *Anopheles* ITS2 sequences have been submitted to NCBI and are available online under GenBank accession numbers XXX.

### *Plasmodium* species identification in *Anopheles* mosquitoes

The following experiments were performed on a LightCycler® 96 Instrument (Roche Diagnostics GmbH, Mannheim, Germany). Initial screening for *Plasmodium spp.* was performed by qPCR following the protocol described by Mangold *et al.* [38]. The PCR reactions were completed with 1 µl of DNA template in total reaction volume of 10 µl. A set of consensus primers was used to amplify a species-specific region of the multicopy 18S rRNA gene (Table 2), along with Euromedex 5X Hot Pol EvaGreen qPCR Mix Plus (ROX), at final concentration of 0.6 µM and 1X, respectively. The thermocycling protocol was 95°C for 12 min, followed by 45 amplification cycles (denaturation at 95°C for 10 s, annealing at 50°C for 10 s and elongation at 72°C for 20 s). A melting-curve analysis was performed at the end of each run by increasing the temperature from 60 °C to 97 °C with continuous fluorescence acquisition to confirm amplicon specificity. Reference melting temperature (Tm) ranges for each *Plasmodium* species were: *P. malariae =* [73.5-75.5°C], *P. falciparum =* [75.5-77.5°C], *P. ovale =* [77.5-79.5°C], and *P. vivax =* [79.5-81.0°C]. Plasmid controls containing partial species-specific 18S rRNA gene sequences were obtained through BEI Resources, NIAID, Bethesda, MD, USA.

**Table 2.**
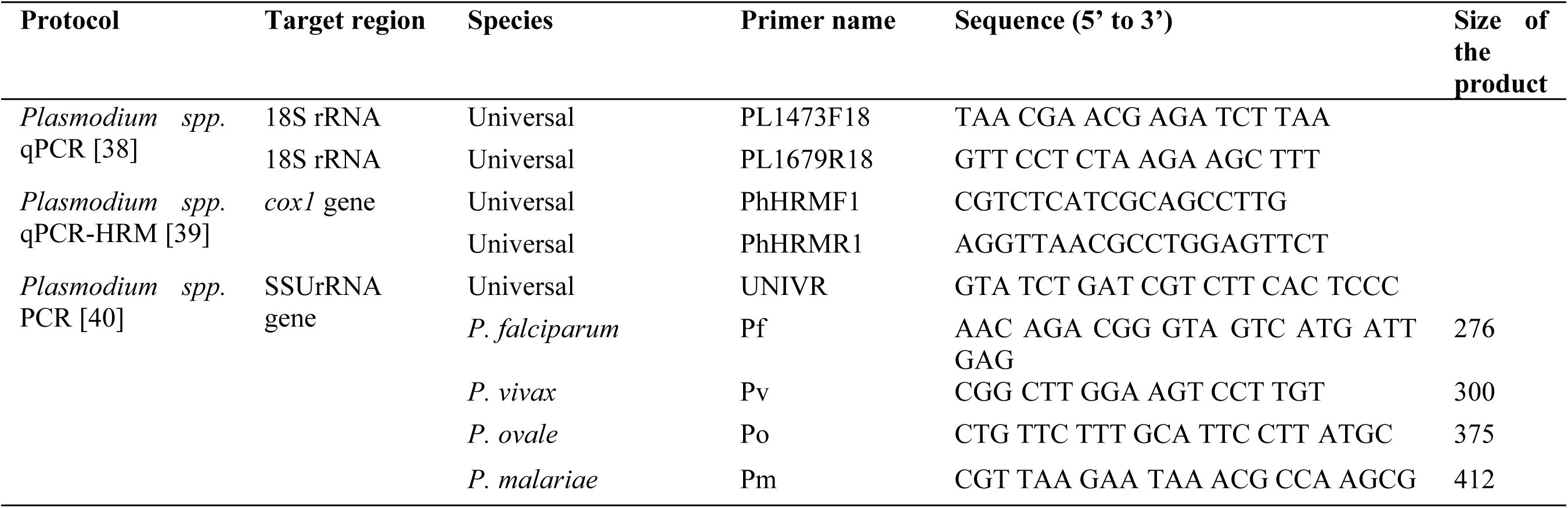
Primers used for molecular detection and identification of *Plasmodium* species in *Anopheles* mosquitoes.

Putatively positive samples according to the screening assay were subsequently analysed using a qPCR High-Resolution Melting (qPCR HRM) protocol adapted from Lamien-Meda *et al.* [39] to discriminate *P. malariae* from *P. falciparum* and to identify mixed-infections. The qPCR HRM assay targeted the mitochondrial *cox1* gene using consensus primers (Table 2) at a final concentration of 0.6 µM each, with Euromedex 5X HotPol EvaGreen HRM Mix (ROX) used at a final concentration of 1X in a 10 µl reaction volume containing 1 µl of template DNA. Cycling consisted of 95 °C for 12 min; 45 cycles of 95 °C for 10 s and 62 °C for 30 s. Fluorescence data were collected continuously at a ramp rate of 0.04 °C/s between 60 °C and 95 °C with 25 acquisitions per °C. Reference Tm values were: *P. malariae =* 77.25 ± 0.03°C, *P. vivax =* 78.11 ± 0.12°C, *P. ovale wallikeri* = 78.53 ± 0.03°C, *P. ovale curtisi* = 78.73 ± 0.05 °C and *P. falciparum* = 79.01 ± 0.12°C. All runs included no-template controls and positive controls for each *Plasmodium* species; data were analysed using the LightCycler® 96 software (version 1.1), and HRM profiles were confirmed by comparison with reference controls.

To go further on the *Plasmodium* identification and prior to sequencing, a PCR targeting 18S Small SubUnit of ribosomal DNA (SSU rDNA) region, developed by Padley *et al.* [40] and commonly used to detect the four major species of human *Plasmodium*, was performed in simplex in order to minimize preferential amplification of the dominant *Plasmodium* species and improve the detection of mixed infections. The PCR reaction mix composition and thermal cycling parameters were identical to those described by Padley *et al.*, with the following modifications: the use of 2.5 U GoTaq G2 Flexi DNA Polymerase (Promega, Madison, WI, USA) and 1X GoTaq Flexi Buffer (Promega, Madison, WI, USA) (Table 2). The PCR products were subjected to electrophoresis on a 2% agarose gel. Lengths of amplified species-specific products are presented in Table 2. Size verified products were commercially sequenced by GENEWIZ Germany GmbH (Leipzig, Germany) using Sanger method with the universal reverse primer. Sequence alignments were then performed within the GenBank dataset (http://www.ncbi.nlm.nih.gov) using the basic local alignment search tool BLASTN algorithm (BLAST, Bethesda, MD, USA). Verified *Plasmodium* 18S sequences have been submitted to NCBI and are available online under GenBank accession numbers XXX.

### Statistical analysis

Accuracy between morphological and molecular identifications was calculated as the proportion of specimens correctly identified by morphology relative to molecular results. Additionally, a partial identification accuracy was calculated for specimens assigned to the same group or complex as the molecularly identified species. Differences in total *Anopheles* abundance between the dry season (July 2024) and the rainy season (December 2024) were assessed using a Wilcoxon matched-pairs signed-rank test. Each sampling night within a village constituted a paired observation (2 nights × 4 villages = 8 pairs). Differences in *Anopheles* species composition between the dry and rainy seasons were assessed using a Freeman-Halton extension of Fisher’s exact test, with p-values estimated by Monte Carlo simulation (10 000 replicates). Species with low observed counts (<5 specimens) were grouped together under “Other *spp.*” category prior to analysis. Differences in the indoor/outdoor distribution of *Anopheles* species were assessed using Pearson’s chi-squared test on contingency tables combining species and collection setting, separately for each season and trap type.

Statistical analyses were performed using R version 4.6.0 (2026-04-24 ucrt). Differences in the indoor/outdoor distribution of Anopheles species were assessed using Pearson’s chi-squared test on contingency tables combining species and collection setting (indoor/outdoor), separately for each season and trap type.

The *Plasmodium* infection rate (IR%) was calculated as the percentage of positive *Anopheles* (head+thorax): IR = number of *Anopheles* positive/total *Anopheles* tested X 100. Differences in sporozoite infection rates between the dry and rainy seasons were analysed using Fisher’s exact test, with odds ratios (OR) and 95% confidence intervals were calculated using the Baptista-Pike method (GraphPad Prism 8.0)

## Results

### *Anopheles* abundance and diversity

A total of 1,811 female mosquitoes were collected across the three sampling campaigns. Of these, 1,157 (63.9%) belonged to the *Anopheles* genus (Table 3), 650 were *Culex* (35.8%), and 4 *Aedes* (0.2%). No *Anopheles* specimens were collected during the daytime capture (between 06:00 and 18:00) with the BGP (20 traps) set up in December 2024. Of all the *Anopheles* females, 1,155 (99.9%) were morphologically identified. The *Anopheles* identified as main vectors, notably *An. gambiae s.l.* and *An. funestus s.l.,* as well as secondary vectors such as *An. marshallii*, *An. coustani*, *An. ziemanni*, *An. paludis*, *An. rufipes*, *An. nili*, *An. pharoensis,* and *An. hancocki*, accounting for 828 (71.6%) of all *Anopheles* specimens collected, were subjected to molecular testing (Table 4). The PCR-RFLP of Fanello *et al.* failed to identify 15 *An. gambiae s.l.* specimens out of 43 (34.9%), while the multiplex PCR by Garros *et al* failed on 47 *An. funestus s.l.* specimens out of 536 (8.8%) (S1 Table). These 62 specimens were therefore subjected to sequencing. The PCR-RFLP successfully differentiated *An. gambiae* from *An. arabiensis*, and results were validated by sequencing. Considering all molecular detection methods, *An. arabiensis* represented one fourth (27,5%) of *An. gambiae* complex collected (11/40), compared to *An. gambiae* representing 70% (28/40) and *An. coluzzii* 2.5% (1/40) (Table 3). From the sequencing of 347 *Anopheles*, 14 species were identified (Table 3). Specimens that showed no reliable match in the GenBank (n=3) or could not be assigned to a named species (n=2) have been regrouped under “unknown *spp.*” (Table 3, Fig. 2). Four species (*An. demeilloni*, *An. implexus, An. nili,* and *An. obscurus*) were identified based solely on morphology. Furthermore, a subset of less common species within the *An. funestus* group, namely *An. leesoni*, *An. parensis*, *An. rivulorum,* and *An. vaneedeni*, was identified using the PCR method described by Garros *et al.* (S1 Table), but were not validated by sequencing. Moreover, most of the specimens in this subset did not belong to the *An. funestus* group, except for four mosquitoes identified as *An. longipalpis* types A (n=3) and C (n=1) (Table 3). Thus, specimens that were not sequenced and not identified as *An*. *funestus* based on the PCR results were considered as “unresolved identification” (Table 3). *An. funestus* was the most abundant species across all surveys, representing 43.4% of all *Anopheles* collected (502/1,157) (Table 3). Focussing on well documented secondary vectors, *An. coustani* was the most abundant (85/1,157, 7.3%), followed by *An. rufipes* (59/1,157, 5.1%), *An. marshallii* (53/1,157, 4.6%), and *An. squamosus* (39/1,157, 3.4%).

**Figure 2:**
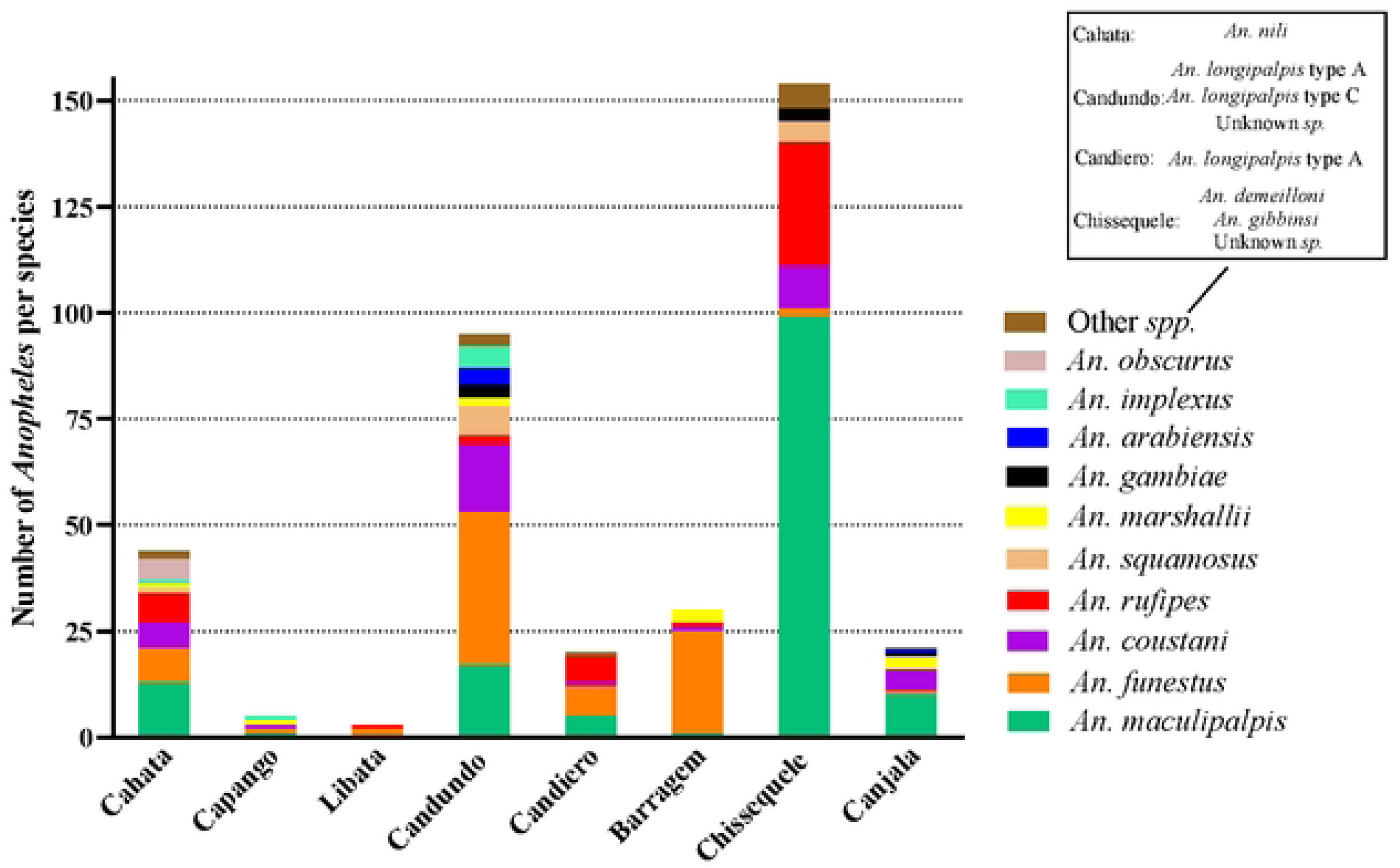
Species composition and abundance of *Anopheles* mosquitoes collected across the eight study villages in the Balombo region, Benguela Province, Angola.

**Table 3:** Number and percentage of *Anopheles* females collected in December 2023, and July, December 2024.

| Species | December 2023 |  | July 2024 |  | December 2024 |  | Total |  |
| --- | --- | --- | --- | --- | --- | --- | --- | --- |
|  | n | % | n | % | n | % | n | % |
| *An. arabiensis* | 5 | 1.3 | - | - | 6 | 1.0 | 11 | 1.0 |
| *An. brunnipes* | - | - | 1 | 0.5 | - | - | 1 | 0.1 |
| *An. coluzzii* | - | - | - | - | 1 | 0.2 | 1 | 0.1 |
| *An. coustani* | 40 | 10.2 | 4 | 2.0 | 41 | 7.3 | 85 | 7.3 |
| *An. demeilloni* | 4 | 1.0 | - | - | - | - | 4 | 0.3 |
| *An. funestus* | 81 (2<sup>a</sup>) | 20.7 | 118 | 60.2 | 303 | 53.2 | 502 | 43.4 |
| *An. gambiae* | 7 | 1.8 | - | - | 21 | 3.7 | 28 | 2.4 |
| *An. gibbinsi* | 1 | 0.2 | - | - | - | - | 1 | 0.1 |
| *An. implexus* | 7 | 1.8 | - | - | - | - | 7 | 0.6 |
| *An. longipalpis* type A | 2 | 0.6 | 1<sup>a</sup> | 0.5 | - | - | 3 | 0.3 |
| *An. longipalpis* type C | 1 | 0.2 | - | - | - | - | 1 | 0.1 |
| *An. maculipalpis* | 146 | 37.1 | 23 | 11.7 | 75 | 13.2 | 244 | 21.1 |
| *An. marshallii* | 10 | 2.5 | 23 | 11.7 | 20 | 3.5 | 53 | 4.6 |
| *An. nili* | 2 | 0.6 | - | - | - | - | 2 | 0.2 |
| *An. obscurus* | 5 | 1.3 | - | - | 1 | 0.2 | 6 | 0.5 |
| *An. rhodesiensis* | - | - | 18 | 9.2 | 54 | 9.5 | 72 | 6.2 |
| *An. rufipes* | 46 | 11.4 | 2 | 1.0 | 11 | 1.9 | 59 | 5.1 |
| *An. squamosus* | 13 | 3.3 | - | - | 26 | 4.6 | 39 | 3.4 |
| Unknown spp. | 2 | 0.5 | - | - | 3 | 0.5 | 7 | 0.4 |
| Unresolved identification | 20 | 5.1 | 6 | 3.2 | 7 | 1.2 | 33 | 2.9 |
| Total | 392 | 100 | 196 | 100 | 569 | 100 | 1157 | 100 |

<sup>a</sup> : Prokopak trap

**Table 4:** Accuracy of morphological identification of *Anopheles* species validated by molecular analysis.

| Morphological identification |  | Molecular identification |  |  |  |  |  |  |  |  |  |  |  |  |  |  |  |  |  |
| --- | --- | --- | --- | --- | --- | --- | --- | --- | --- | --- | --- | --- | --- | --- | --- | --- | --- | --- | --- |
| Species | n (total) | *An. funestus* | *An. longipalpis* type A | *An. longipalpis* type C | *An. gambiae* | *An. coluzzii* | *An. arabiensis* | *An. marshalli* i | *An. squamosus* | *An. rufipes* | *An. maculipalpis* | *An. coustani* | *An. rhodesiensis* | *An. brunnipes* | *An. gibbinsi* | Unknown spp. | Unresolved identification | Identification accuracy (%) | Complex or group identification accuracy (%) |
| *An. funestus s.l.* | 574 | 492 (444<sup>a</sup>) | 3 | 1 | 2 | 1 | - | 17 | 11 | 9 | 5 | 2 | 1 | 1 | - | 1 | (28<sup>a</sup>) | 85.7 | 86.4 |
| *An. gambiae s.l.* | 43 | 1 | - | - | 24 (20<sup>a</sup>) | - | 11 (9<sup>a</sup>) | - | - | 3 | - | - | - | - | - | 1 | 3 | 55.8 | 81.4 |
| *An. marshallii* | 46 (53) | 3 | - | - | - | - | - | 30 | - | 11 | 1 | 1 | - | - | - | - | - | 65.2 | 65.2 |
| *An. coustani* | 19 (20) | - | - | - | - | - | - | - | 7 | - | - | 12 | - | - | - | - | - | 63.2 | 63.1 |
| *An. ziemanni* | 78 (85) | 2 | - | - | 1 | - | - | - | 16 | 2 | 7 | 49 | - | - | - | 1 | - | 0 | 62.8 |
| *An. paludis* | 17 | - | - | - | - | - | - | - | 3 | - | 4 | 10 | - | - | - | - | - | 0 | 58.8 |
| *An. rufipes* | 44 (52) | 4 | - | - | 1 | - | - | 1 | 2 | 22 | 12 | 1 | - | - | - | 1 | - | 50.0 | 50.0 |
| *An. nili* | 1 (3) | - | - | - | - | - | - | - | - | - | - | - | 1 | - | - | - | - | 0 | 0 |
| *An. pharoensis* | 2 (2) | - | - | - | - | - | - | - | - | - | - | - | - | - | 1 | 1 | - | 0 | 0 |
| *An. hancocki* | 4 (4) | - | - | - | - | - | - | - | - | 4 | - | - | - | - | - | - | - | 0 | 0 |
| Total | 828 | 502 | 3 | 1 | 28 | 1 | 11 | 48 | 39 | 51 | 29 | 75 | 2 | 1 | 1 | 5 | 31 |  |  |

<sup>a</sup> : PCR results.

### Identification accuracy

When comparing the morphological and molecular species identifications, *An. funestus* had the highest accuracy with 85.7% (492/574), followed by *An. marshallii* with 65.2% (30/46), *An. coustani* with 63.2% (12/19), *An. gambiae* with 55.8% (24/43), and *An. rufipes* with 50% (22/44) (Table 4). When comparing morphological to molecular identification of specimens belonging to the same group or complex, the accuracy reached 81.4% (35/43) for the *An. gambiae* complex. Morphological identification failed on *An. nili*, *An. paludis, An. pharoensis, An. hancocki,* and *An. ziemanni*. However, considering partial identification of *An. paludis* and *An. ziemanni* as member of *An. coustani* group, the accuracy at the group level reached 58.8% (10/17) and 62.8% (50/78), respectively.

### Preliminary campaign and village selection

In December 2023, 392 female *Anopheles* were collected (S2 Table). A total of 15 *Anopheles* species were identified in this entomological survey (11 molecularly and 4 morphologically), while 22 specimens (5.6%) remain unidentified (unknown *spp*. and unresolved identification). *An. maculipalpis* was the most abundant species (37.2%, 146/392), followed by *An. funestus* (20.6%, 81/392) (Fig. 2). Other main malaria vectors like *An. gambiae* (1.8%, 7/392) and *An. arabiensis* (1.3%, 5/393) were also captured. Secondary vectors like *An. rufipes* (11.7%, 46/392), *An. coustani* (10.2%, 40/392), and to a less extent *An. squamosus* (3.3%, 13/392) and *An. marshallii* (2.5%, 10/392) were also collected. When considering the distribution of *Anopheles* specimens per village, nearly 80% (310/393) of the collected *Anopheles* were obtained from three villages, Cahata (n=48), Candundo (n=96), and Chissequele (n=166) (Fig. 2). A fourth village, Libata, was also selected for the 2024 sampling campaigns due to the significant experience gained in that village as part of the Balombo project, during which mosquito abundance and *Anopheles* species diversity were documented.

### Vector seasonality

Comparisons between both seasons of 2024 were conducted in the four selected villages (Table 5 and Fig. 3). *Anopheles* abundance was significantly higher during the rainy season (n=569) than during the dry season (n=196) across all villages (exact two-tailed Wilcoxon matched-pairs signed-rank test: W = 36, *P* = 0.0078, median difference = 20.5). Candundo showed the highest *Anopheles* abundance across both seasons (dry: 127; rainy: 344), while Chissequele and Libata yielded the lowest catches (dry: 9 and 6; rainy: 32 and 47, respectively). A total of eight *Anopheles* species were collected during the dry season, while 11 *Anopheles* species were identified during the rainy season. Considering all villages combined, *Anopheles* species composition differed significantly between the dry and rainy seasons (Fisher’s exact test, Monte Carlo simulation with 10 000 replicates, *P* < 0.0001), as well as in Cahata (*P* < 0.0001) and Candundo (*P* = 0.0074) when analysed individually. No significant differences were detected in Chissequele (*P* = 0.339) or Libata (*P* = 0.180), likely due to the limited number of specimens collected during the dry season at these sites (<10 specimens at both locations). *An. funestus* had the highest relative proportion in both dry (60.2%, 118/196) and rainy seasons (53.2%, 303/569) followed by *An. maculipalpis* (dry: 11.7%; rainy: 13.2%). It should be noted that the dominance of *An. funestus* within the anopheline populations was less pronounced in Cahata, Chissequele, and Libata than in Candundo during both seasons (Table 5). Consequently, the high overall relative proportion of this species was largely driven by the large number of *An. funestus* specimens collected in Candundo (79.3%, 334/421). The primary vectors of the *An. gambiae* complex (*An. arabiensis*, *An. coluzzii,* and *An. gambiae*) were collected only during the rainy season, as was the secondary vector *An. squamosus*. Other secondary vectors (*An. coustani*, *An. marshallii,* and *An. rufipes*) were collected during both seasons, with relatively higher proportions during the rainy season, except for *An. marshallii* (Table 5).

**Figure 3:**
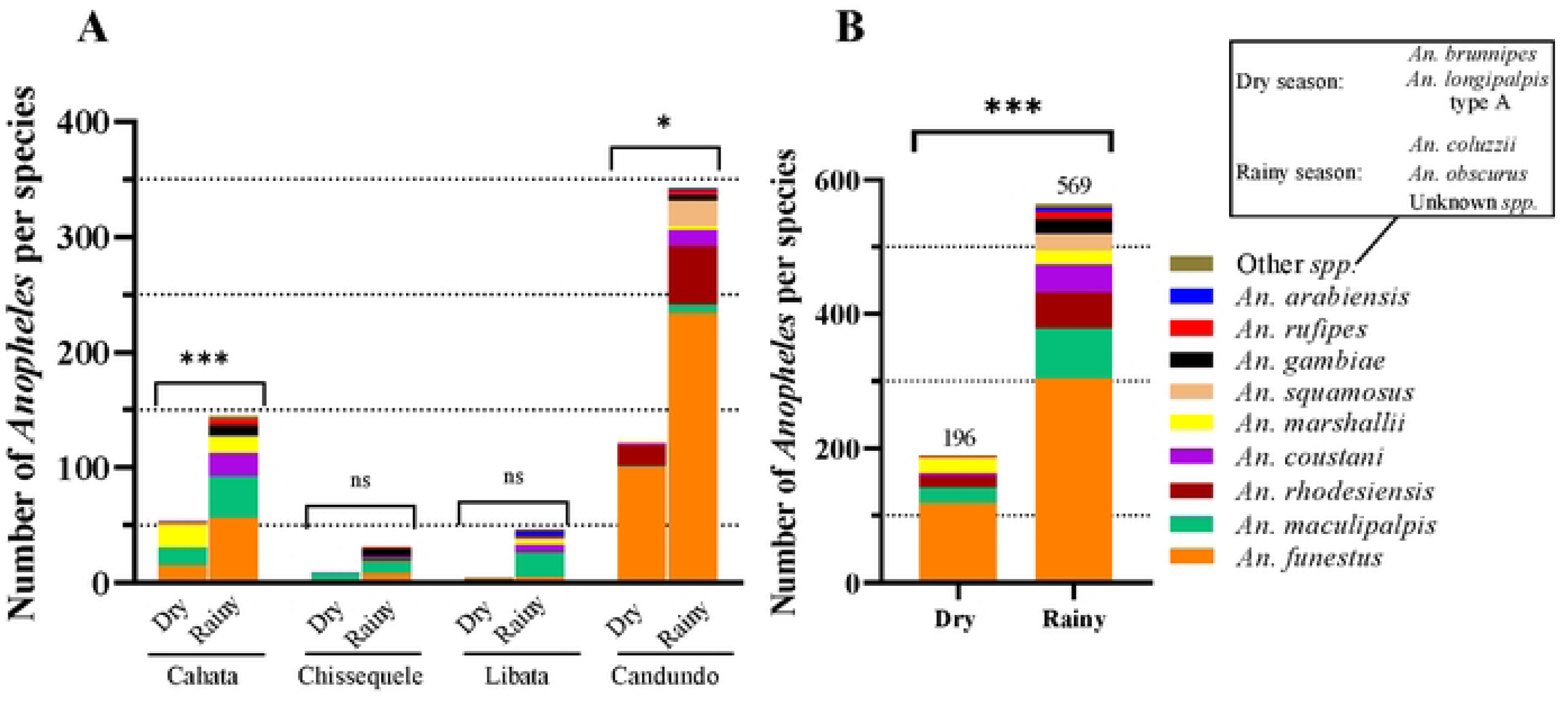
Differences in *Anopheles* species composition by seasons and across villages (A) by season only (B) in Balombo, 2024 (Fisher’s exact test).

**Table 5:**
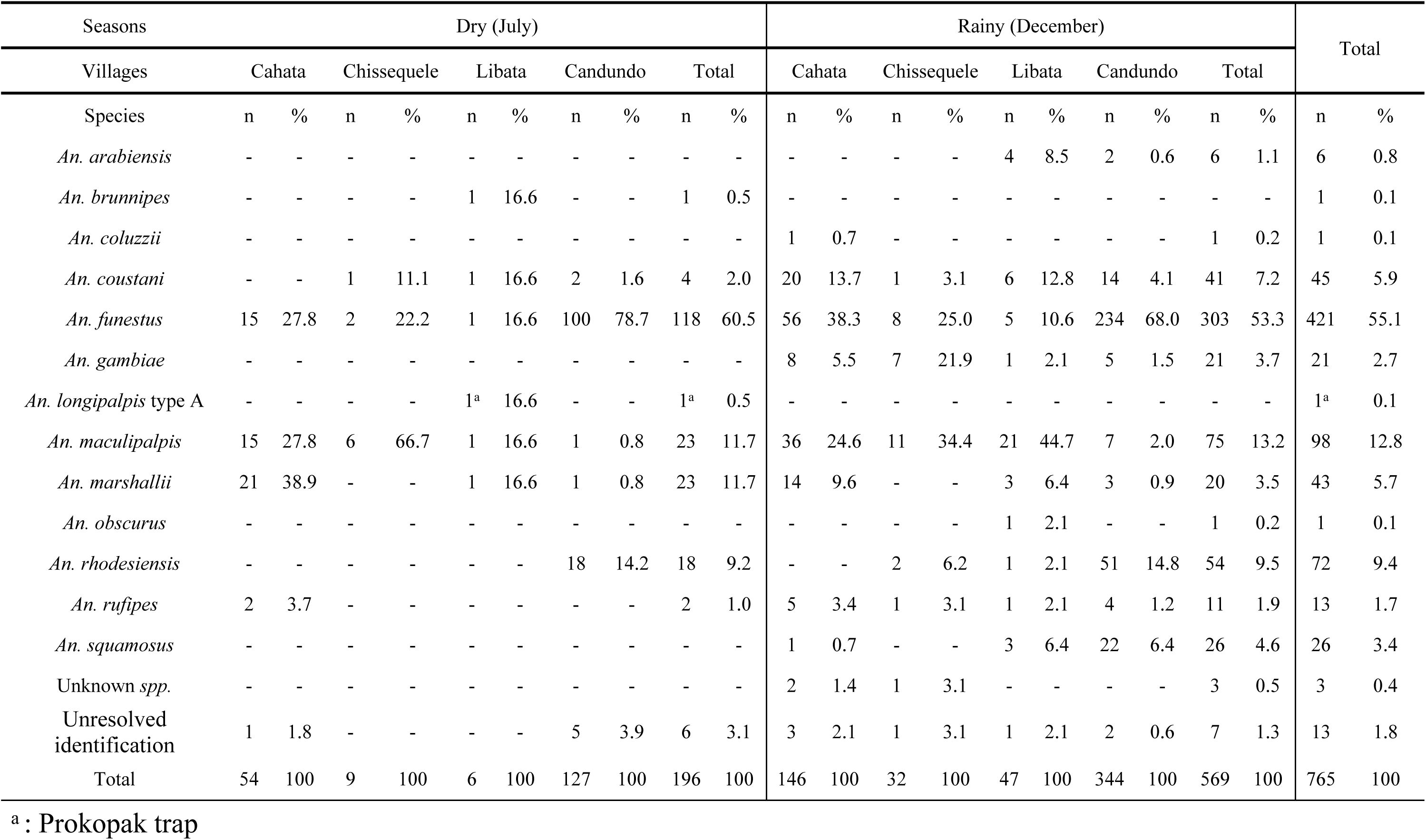
Abundance and diversity of *Anopheles* collected in 4 villages of Balombo in dry (July) and rainy (December) seasons in 2024.

### Indoor and outdoor host-seeking activities of *Anopheles* mosquitoes

Indoor and outdoor mosquito collections were compared separately for each trap type (CDC-LT and BGP) within each season (Fig. 4A), and separately across seasons regardless of trap type (Fig. 4B). The Prokopak collection conducted in July 2024 was excluded from the analysis because only one female *Anopheles* was captured. During both seasons, indoor traps collected more *Anopheles* specimens (dry: 117; rainy: 383) than outdoor ones (dry: 78, rainy: 186) (S3 Table). While more *Anopheles* species were collected outside (n=7) than inside (n=5) during the dry season, and more species were collected indoor (n=12) than outdoor (n=9) during the rainy season (S3 Table). Due to low sample sizes for several species, analyses were restricted to species with at least 20 specimens. Regardless of the villages and considering each trap type independently during the dry and the rainy season, the indoor/outdoor distribution of *Anopheles* species were highly significantly different (CDC dry season: χ² = 53.71, df = 3, *P* < 0.0001; BGP dry season: χ² = 40.8, df = 3, *P* < 0.0001; CDC rainy season: χ² = 188.4, df = 8, *P* < 0.0001; BGP rainy season: χ² = 113.7, df = 6, *P* < 0.0001) (Fig. 4A). Pooling trap types did not change this pattern: the indoor/outdoor distribution remained significantly heterogeneous in both seasons (dry season: χ² = 93.17, df = 4, *P* < 0.0001; rainy season: χ² = 282.8, df = 9, *P* < 0.0001) (Fig. 4B). *An. funestus* was predominantly collected indoors during both seasons in Candundo, while the proportion of indoor captures was lower in Cahata, Chissequele and Libata (data not shown). Low sample sizes for other primary vector species within the *Anopheles gambiae* complex limited further analysis; however, *An. gambiae* was predominantly collected indoors and only during the rainy season. In contrast, during the rainy season, patterns were less consistent among secondary vectors. *An. coustani* was mainly collected outdoors, whereas *An. squamosus* was captured in similar proportions regardless of indoor or outdoor sampling location. *An. marshallii* showed a seasonal shift, being more frequently collected outdoors during the dry season and indoors during the rainy season (Fig. 4A and 4B). Zoophilic *Anopheles* species like *An. maculipalpis* or *An. rhodesiensis*, were mainly collected outside in both seasons.

**Figure 4:**
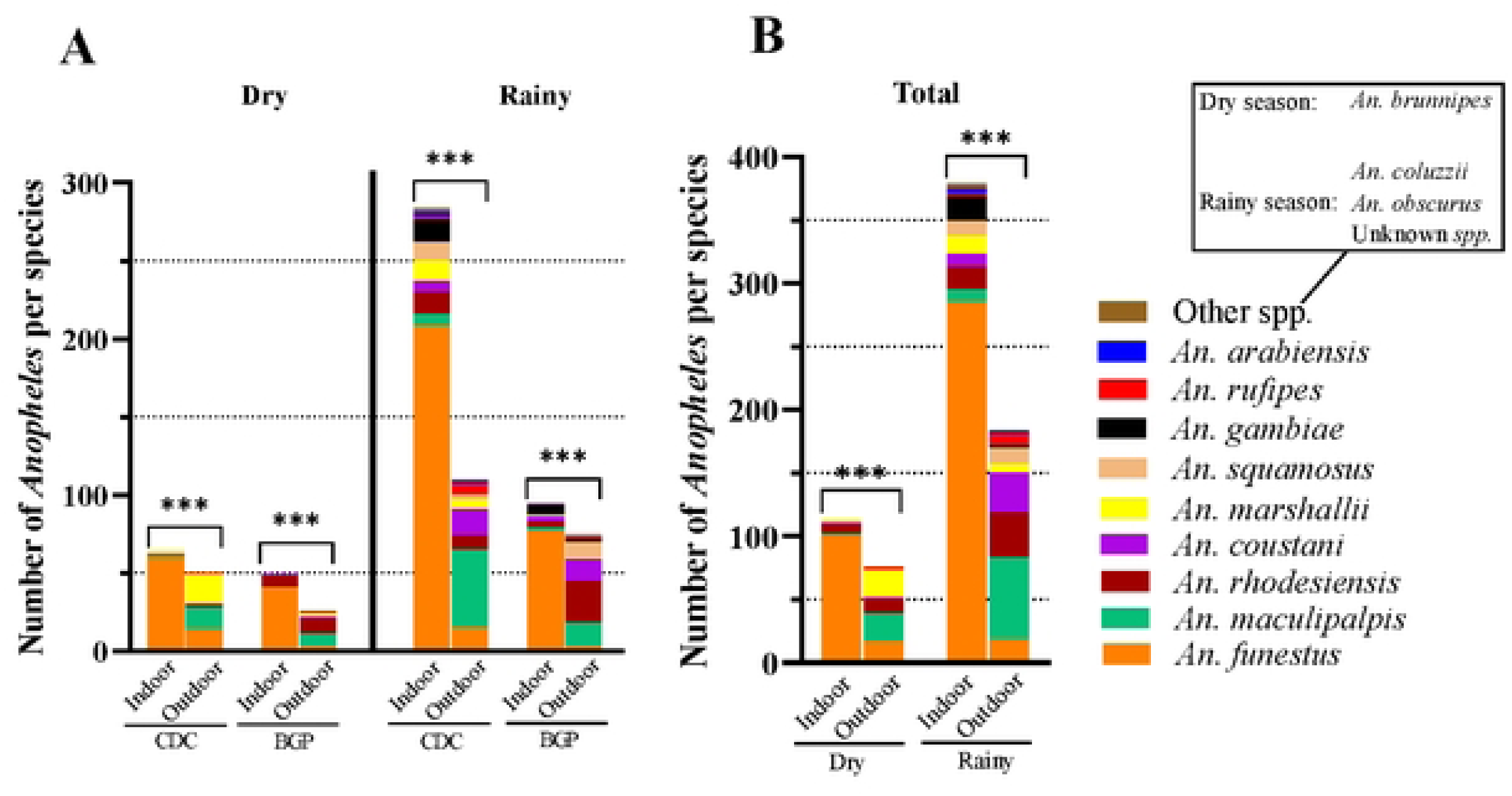
Differences in *Anopheles* species composition between indoor and outdoor collections, by trap type and season (A), and by season only (B) in Balombo in 2024 (χ²).

### Comparison of trapping methods

Overall, in 2024, CDC-LT accounted for 64.8% (n=750) of collected *Anopheles*, while BGP contributed to 35.2% (n=408) and Prokopak to only 0.1% (n=1) (Addition file 3). Consequently, the comparison between the mosquito samples collected by CDC-LT and by BGP was conducted without considering the prokopak collection. During both seasons, the mean number of *Anopheles* per trap per night was higher with CDC-LT (dry: 2.4; rainy: 8.3) compared to BGP (dry: 1.6; rainy: 3.5). No difference was observed in the *Anopheles* species composition collected by both types of traps in both seasons, with few exceptions on rare species with low sample sizes (n=1), such as *An. coluzzii* only collected with the CDC-LT, and *An. brunnipes* and *An. obscurus* with BGP.

### Detection of *Plasmodium spp*. DNA in *Anopheles* mosquitoes

Head-thoraces of the 1,085 *Anophele*s from the three campaigns were subjected to molecular analyses to detect *Plasmodium* infection. Among the 840 specimens morphologically identified as main or secondary vectors and screened with qPCR, 65 were selected based on their melting peak profile for qPCR HRM for better discrimination between *Plasmodium* species (Table 6). Following the unexpected detection of *P. falciparum* in a specimen of *An. maculipalpis*, a total of 245 *An. maculipalpis* were subsequently tested using qPCR HRM. Amplification of the 18S gene of *Plasmodium* was also done separately for *P. falciparum*, *P. malariae,* and *P. ovale* on a subset of 78 *Anopheles* specimens, and sequencing was performed on 48 specimens. No other specimens of *An. maculipalpis* tested positive.

**Table 6:** Summary of positive *Anopheles* females collected across the sampling campaign of Balombo between 2023 and 2024, based on different detection methods.

| Detection methods |  | qPCR 18S<br>(Mangold *et al*) | qPCR HRM<br>*coxI* (Lamien-Meda *et al*) | PCR 18S<br>(Padley *et al*) | Sequencing 18S |
| --- | --- | --- | --- | --- | --- |
| Sampling campaign | Species | n=840 | n=310 | n=78 | n=48 |
| December 2023 | *P. falciparum* | 3 | 3 | 4 | 4 |
|  | *P. malariae* | 2 | 2* | 1 | 1 |
|  | Total | 5 | 5* | 5 | 5 |
| July 2024 | *P. falciparum* | 0 | 2 | 3 | 3 |
|  | *P. malariae* | 3 | 0 | 0 | 0 |
|  | *Pf + Pm* | 0 | 1 | 0 | 0 |
|  | Total | 3 | 3 | 3 | 3 |
| December 2024 | *P. falciparum* | 22 | 23 | 21 | 20 |
|  | *P. malariae* | 1 | 3 | 2 | 1 |
|  | *Pf + Pm* | 4 | 1 | 1 | 1 |
|  | Total | 27 | 27 | 24 | 22 |
| Total |  | 35 | 35 | 32 | 30 |

\*: *An. maculipalpis* detected positive to *P. malariae* in qPCR HRM but to *P. falciparum* in PCR and sequencing

In total, 35 *Anopheles* specimens were tested positive for *Plasmodium spp.* based on both qPCR 18S-Mangold and HRM analysis. Among them, 29 (82.9%) were positive for *P. falciparum*, 4 (11.4%) for *P. malariae,* and 2 (5.7%) were co-infected with both species (Table 7). Conventional PCR successfully detected *Plasmodium spp.* in 32 *Anopheles* and failed to detect infection in 3 high-Cq positive specimens (Cq between 34-35), which were found using both qPCR (Mangold and HRM) (Table 6). Most positive *Anopheles* were collected during the rainy season, with 5 in December 2023 and 27 in December 2024 (91.4%) compared to only 3 (8.6%) in July 2024 (dry season) (Table 7). Similarly, a greater diversity of *Anopheles* species infected with *Plasmodium* was observed during the rainy season (n=6) than during the dry season (n=2) (Table 7).

**Table 7:** *Plasmodium* infection rate in *Anopheles* species collected across sampling campaigns between 2023 and 2024.

| Sampling surveys | Species | n | Villages | *P. falciparum* | *P. malariae* | *Pf + Pm* | IR (%) |
| --- | --- | --- | --- | --- | --- | --- | --- |
| December 2023 | *An. funestus* | 81 | Candundo | 1 | 1 | - | 4.9 |
|  |  |  | Barragem | 2 | - | - |  |
|  | *An. maculipalpis* | 146 | Candundo | 1 | - | - | 0.7 |
| July 2024 | *An. funestus* | 118 | Candundo | 2 | - | - | 1.7 |
|  | *An. marshallii* | 20 | Cahata | - | - | 1 | 5 |
| December 2024 | *An. funestus* | 223 | Candundo | 13 | - | - | 9.4 |
|  |  |  | Cahata | 5 | 2 | - |  |
|  |  |  | Libata | 1 | - | - |  |
|  | *An. marshallii* | 21 | Cahata | 2 | - | 1 | 14.3 |
|  | *An. gambiae* | 22 | Chissequele | 1 | - | - | 4.5 |
|  | *An. arabiensis* | 6 | Libata | - | 1 | - | 16.7 |
|  | *An. rufipes* | 11 | Chissequele | 1 | - | - | 9.1 |
|  | Total |  |  | 29 | 4 | 2 | - |

*An. funestus* was the most represented species among positive mosquitoes (77.1%, 27/35), collected across all surveys. It was followed by *An. marshallii* (11.4%, 4/35), collected during both seasons in 2024. Four other species, *An. arabiensis*, *An. gambiae*, *An. maculipalpis,* and *An. rufipes*, were each detected as positive only once (2.9%, 1/35). Candundo accounted for the highest proportion of positive specimens (51.4%, 18/35), followed by Cahata (31.4%, 11/35) and Barragem, Chissequele, Libata (5.7% each, 2/35).

The overall sporozoite infection rate (IR) was significantly higher during the rainy season than during the dry season (9.54% vs. 2.17%; Fisher’s exact test, OR = 4.75, 95% CI: 1.52–15.09, *P* = 0.0044). When the analysis was limited to *An. funestus*, the only species with sufficient sample size during both seasons (n= 118 and 223 respectively), the seasonal difference remained significant (IR: 1.69% in dry season vs. 9.42% in rainy season; Fisher’s exact test, OR = 6.03, 95% CI: 1.56–26.4, *P* = 0.0057). This result indicates that *An. funestus* was approximately six times more likely to be infected during the rainy season than during the dry season in 2024.

## Discussion

This study represents the first comprehensive entomological survey conducted in the Balombo region in 17 years, providing updated insights into the *Anopheles* vectors after nearly two decades of vector control interventions, mainly based on the distribution of long-lasting insecticidal nets (Réf 24). The results highlight the diversity of *Anopheles* species in villages of Balombo during both the dry and rainy seasons and report, for the first time, the detection of *P. malariae* in *Anopheles* mosquitoes in the Benguela Province. Moreover, the main vector, *An. funestus,* is still widespread [25] and was found to be infected during both seasons. In addition, the study also reveals for the first time that, in the Balombo region, five other *Anopheles* species were infected with *P. falciparum* or *P. malariae*, including cases of coinfection, suggesting a potential change in the vectorial situation and highlighting the complexity of local malaria transmission dynamics. The study aimed to investigate the entomological factors that may underlie the epidemiological shift observed in the villages of Balombo since 2016. Annual field reports from the Sonamet MCP have documented an increasing malaria prevalence among villagers in the Balombo region over the years, together with the identification of non-*falciparum* malaria cases, mainly *P. malariae*, but also occasionally *P. ovale,* and rare *P. vivax* infections [29]. Changes in the vector population, including behavioural adaptation, and the potential involvement of secondary vector species, were therefore considered among the possible explanations. Our results show that *An. funestus* remained the most abundant species, accounting for 43% of the collected specimens, consistent with the findings of the last major entomological study [24], in which this major vector species represented 50% of the collected specimens. This observation is also supported by ecological observations, as suitable larval habitats for this species were identified near each village across the different surveys [41]. Indeed, all villages of Balombo are located close to a stream or a river (<1km), allowing permanent water bodies with emergent vegetation to persist even during the dry season [5]. These environmental conditions may partly explain the sustainability of malaria transmission observed throughout the year in the Balombo villages [5,42,43]. The results also showed a high diversity of *Anopheles* species in the studied sites, with a total of 18 species collected during both seasons. Beside *An. funestus*, three additional major malaria vector species were identified, all belonging to the *An. gambiae* complex, *An. arabiensis*, *An. coluzzii,* and *An. gambiae*, which is consistent with previous reports from Angola and Benguela Province [23–25,44]. *Anopheles* species, commonly described in the literature as secondary malaria vectors [18,19], were collected and molecularly identified as *An. coustani*, *An. marshallii*, *An. rufipes*, and *An. squamosus*, which is also consistent with previous studies conducted in Angola [21,25]. Furthermore, specimens collected in December 2023 and morphologically identified as *An. nili* were excluded from the analyses, because the only sequenced specimen was molecularly identified as *An. rhodesiensis*, even though *An. nili* is known to occur in Angola [21]. Several zoophagic species (n=8), previously reported in the Angolan anopheline fauna [19], were also collected, including *An. brunnipes*, *An. demeilloni*, *An. longipalpis* types A and C, *An. implexus*, *An. obscurus*, *An. maculipalpis,* and *An. rhodesiensis*. This study reports the first detection of *An. gibbinsi* in Angola, although this species has already been recorded in Central and East Africa [19], where it is considered a primarily zoophilic or opportunistic species exhibiting exophagic and exophilic behaviour. It was collected using an outdoor CDC-LT in Chissequele during the first rainy season of December 2023. Cases of *P. falciparum* infection have occasionally been observed in *An. demeilloni*, *An. longipalpis* type C, and *An. gibbinsi*, mainly in East Africa [20,45,46], suggesting that their potential role as secondary malaria vectors in Angola warrants further investigation.

The distribution of *Anopheles* species is constrained by environmental suitability [44]. The results of this study suggest that the diversity and abundance of *Anopheles* species decreased by more than half during the dry season (7 species, for a total of 196 specimens collected) compared to the rainy season (18 species, for a total of 569 specimens collected), with marked heterogeneity observed from one village to another. Four species of public health importance were collected exclusively during the rainy season, namely *An. arabiensis*, *An. coluzzii*, *An. gambiae,* and *An. squamosus*. Temperatures affect the development rate and survival of mosquitoes, whereas rainfall influences the presence of larval habitats and, consequently, affects the diversity and the density of *Anopheles* populations [41,42,47]. In general, malaria incidence is lower during the dry season due to the spatial confinement and reduction of *Anopheles* populations. Consequently, many entomological studies have focused primarily on the rainy season, when high malaria transmission peaks typically begin to occur [48].

This study aimed to address the lack of available seasonal data on Anopheles populations in the Benguela Province; however, it should be noted that some interpretations should be viewed with caution due to the relatively small number of specimens collected in some villages (e.g., Chissequele and Libata in July 2024). Environmental conditions, such as heavy rain in December and relatively cold night (<10°C) in July, have affected the capture of *Anopheles*. The performance of CO_2_-dependant traps, such as BGP, is considerably reduced in rainy conditions, and mosquitoes caught in outdoor traps may end up completely crushed and soaked. Thus, the type of trap may also have skewed some observed trends. Although the BGP and CDC-LT appeared complementary in terms of species diversity collected during the December 2023 survey, this was not observed during subsequent surveys. Moreover, the number of *Anopheles* specimens collected was consistently higher with CDC-LT across all surveys, by approximately 1.5-fold during the dry season and up to 4-fold during the rainy season. Due to the very low number of *Anopheles* collected with the Prokopak (n=3), this method was no longer used after July 2024. Its low effectiveness may be explained by the early sunrise (05:30 in December and 05:45 in July) and morning activities of villagers (ventilation of the rooms by opening the windows). Despite the absence of human landing catch (HLC) and the limitations associated with the use of BGP and CDC-LT for studying the *Anopheles* behaviour [49,50], these collecting methods still provide valuable information on vector abundance, vector density, and sporozoite infection rates [51–53].

Accurate identification of the captured *Anopheles* species is crucial for understanding the malaria dynamics, particularly when changes in human parasite prevalence, vector behaviour, or species composition are observed [15]. It also supports the selection of appropriate vector control strategies. In this study, accuracy was considered correct when morphologically identified specimens belonged to the same *Anopheles* group or complex as those identified by molecular methods. By definition, members of a species complex cannot be differentiated using morphological identification keys, and species within a group or subgroup are often difficult to distinguish due to overlapping morphological characters, therefore, reliable species identification must be complemented by molecular approach [37,54]. The results indicate that morphological identification was consistent for the major malaria vectors in Angola, with accuracies reaching 85% for both *An. gambiae s.l.* and *An. funestus*. Although morphological identification accuracy was lower for secondary vectors (ranging from 50% to 65.2%), the overall performance remained acceptable. Several less frequently encountered species, such as *An. brunnipes* and *An. gibbinsi*, were not accurately identified, reflecting the paucity of robust morphological criteria available for the identification of rare *Anopheles* species. These findings are consistent with previous studies reporting high identification accuracy for primary vectors, but lower performance for less common species [55,56]. Nonetheless, the accuracy observed for primary vectors is slightly lower-than-expected, and could be explained by damage of key morphological characters caused by suction through CDC-LT fan blades and by heavy rainfall during collections in Balombo [56,57]. The discordance observed between PCR-based identification and sequencing results for specimens belonging to the *An. funestus* group, may be explained by misinterpretation of PCR amplification bands on agarose gels. Similar issues have previously been reported with *An. longipalpis* type C misidentified as *An. vaneedeni* or *An. parensis* using rDNA ITS region [58], and species of the *An. gambiae* complex being misidentified as *An. leesoni*, a member of the *An. funestus* group [59]. Taken together, these observations combined with the first detection of *An. gibbinsi* in Angola and the collection of specimens for which no corresponding sequences are available in GenBank, underscore the importance of integrating molecular tools into entomological surveillance.

Growing attention has been given to the role of so-called secondary vector species [60–62]. In highly diverse vectorial ecosystems, such as those observed at the study sites, identifying the *Anopheles* species that actively contribute to malaria transmission is the first step toward implementing effective vector control measures. In this study, the first qPCR method targeting the 18S gene [38] lacked specificity, and repeatedly failed to discriminate *P. malariae* from *P. falciparum* in positive specimens, a limitation that was also observed with a conventional multiplex PCR assay targeting the same gene [40]. Similar observations have previously been reported in the literature. According to Demas *et al.* [63], the majority of PCR-based methods still rely on the *Plasmodium* 18S rRNA gene targets, historically considered among the most suitable marker for malaria parasite detection [63]. However, its ability to detect mixed infections in multiplex assays is limited without the use of nested PCR. In contrast, the use of *cox1* gene combined with the qPCR-HRM method developed by Lamien-Meda *et al.* [39], enabled confident detection and discrimination of *Plasmodium* species in 35 *Anopheles* specimens. Most *Anopheles* specimens positive to *Plasmodium spp.* belonged to *An. funestus* (77.1%, 27/35), thereby confirming its role as the primary malaria vector in the Balombo region, consistent with recent findings in Benguela [23], and more globally in Angola [24]. *Anopheles funestus* was found positive to *P. falciparum* in all surveys and to *P. malariae* during both rainy season collections. This study also reports the first detection of *P. malariae* in *Anopheles* mosquitoes in the Benguela Province, although non-falciparum infections in humans have already been reported in this province and elsewhere in Angola [29,64]. *An. marshallii* was the second most frequently infected species in this study (11.4%, 4/35), with specimens tested positive for *P. falciparum* in December 2024 and coinfections with *P. malariae* and *P. falciparum* detected in July and December 2024. Although the role of this species in malaria transmission has previously been reported in Central Africa, this is to our knowledge the first evidence supporting its potential contribution in Angola [18,65]. Finally, the detection of *Plasmodium spp.* in specimens of *An. arabiensis*, *An. gambiae*, *An. maculipalpis,* and *An. rufipes* (2.9% each, 1/35) highlights the complexity of the malaria vector system in the Balombo region, particularly during the rainy season. *An. arabiensis* and *An. gambiae* are the major malaria vector species documented in Angola [44], while it is the first detection of *P. falciparum* in *An. rufipes* in the Benguela Province. The role of *An. rufipes* as a secondary vector in malaria transmission has been documented in several parts of Africa [19,60] and has also been suggested in Angola [21]. Regarding the detection of *P. falciparum* in *An. maculipalpis* in December 2023, its potential involvement in malaria transmission was suggested by Hamon *&* Mouchet in 1961 [66], although more recent studies did not considered it as a public health concern [19,67]. High infection rates were observed in several *Anopheles* species in this study, which may be partly explained by the use of light trap that tend to attract and capture gravid female *Anopheles* and resting mosquitoes, which are detected positives to *Plasmodium* at higher rates [49,50], leading to an overestimation of the infection rate. Moreover, these numbers need to be interpreted with caution since low capture numbers for some species do not allow robust estimation of the infection rates. For instance, one infected *An. arabiensis* among the six specimens tested resulted in an apparent infection rate of 16.7%. A high infection rate for *An. funestus* was also reported elsewhere in Angola [23,24]. Nevertheless, infection rates in both *An. funestus* and *An. marshallii* were higher during the rainy season than the dry season, consistent with previous observations for *An. funestus* in the Benguela Province [23], as higher temperature and higher vector densities tend to be associated with higher infection rates [68,69]. It is noteworthy that *An. funestus* did not exhibit marked endophilic behaviour in the Balombo villages, except in Candundo (during both seasons) and Cahata (only during the rainy season), where the majority of *An. funestus* were captured indoors. However, it is also in these two villages that the largest number of *An. funestus* specimens were recorded. Thus, combining data from all villages may mask local differences in *An. funestus* behaviour because the overall pattern is largely driven by the high abundance of *An. funestus* in Candundo, where the species consistently displayed endophilic/endophagic behaviour. Additional *Anopheles* collections are needed in Balombo to further investigate these observations, particularly in villages where small sample sizes limited the analysis of *An. funestus* exophilic behaviour. Furthermore, conducting monthly collections for at least one year would allow a more robust assessment of *Anopheles* behaviour in relation to seasonal variations.

Several hypotheses may explain the differences observed between Candundo and the other study sites. Candundo is highly isolated compared to the other villages, therefore it was not included in the intensive vector control campaigns conducted by the MCP (Sonamet Company) between 2007 and 2011. During these campaigns, a 81% decrease in *Anopheles* abundance was reported in the six other villages covered by the MCP vector control program between 2008 and 2009 [25]. It is then possible that a selection of individuals with more exophagic/exophilic behaviours may have occurred, alongside with changes in species composition. This last point is supported by the higher proportions of non-funestus species in Chissequele and Cahata compared to Candundo. These observations are consistent with the literature describing the effects of vector control interventions on *Anopheles* populations [11,15,70]. In the case of Balombo, referring to “residual malaria transmission” may be misleading, as these terms refer to transmission that persists after total universal coverage has been achieved through effective vector control [11,71], whereas LLINs coverage in Balombo villages is currently far from universal. However, the detection of *Plasmodium spp.* in secondary vector species in this study may result from the intense vector control program followed by an interruption in LLINs supply to villagers between 2012 and 2016 [72]. Interestingly, *P. malariae* was detected not only in the primary vector *An. funestus* but also in other *Anopheles* species, suggesting that malaria transmission in Balombo may involve a broader mix of vectors than previously reported [25]. Although the contribution of these secondary vectors to transmission remains uncertain, their infection with *Plasmodium spp.* indicates that they may participate in maintaining local transmission dynamics. This observation is particularly relevant given the increasing detection of non-falciparum malaria infections, mainly *P. malariae*, among the human population since 2018 [29]. Together, these findings raise the hypothesis that changes in vector composition, and the involvement of secondary vectors may contribute to the observed epidemiological shift in Balombo. Given the demonstrated presence of infected secondary vectors, which are anthropophilic and predominantly outdoor-biters, it is imperative to integrate complementary interventions targeting outdoor transmission [62].

## Abbreviations

ITNs: Insecticide-Treated Nets
LLINs: Long-Lasting Insecticidal Nets
IRS: Indoor Residual Spraying
NMCP: Angolan National Malaria Control Program
MCP: Malaria Control Program of Sonamet Company
CDC-LT: Centers for Disease Control - Light Trap
BGP: Biogent BG-Pro trap
ITS2: Internal Transcribed Spacer 2
PCR: Polymerase Chain Reaction
RFLP: Restriction Fragment Length Polymorphism
qPCR: Quantitative Polymerase Chain Reaction
HRM: High-Resolution Melting

## Acknowledgments

The authors sincerely thank the populations of Barragem, Cahata, Candiero, Candundo, Canjala, Capango, Chissequele, Libata (Balombo, Angola) for their participation and cooperation throughout this study. We are thankful to the Angolese Sonamet Company, based at Lobito, and its Medical Department for their valuable involvement and logistical support within the framework of the Malaria Control Program (MCP). We also acknowledge the Subsea7 Company, which financially supported the laboratory study conducted at IRD, Montpellier. We further thank the National and Provincial Public Health authorities of Angola for authorizing the field surveys. Finally, we express our sincere gratitude to the French Ambassy in Angola for its confidence in this project, its financial support, and its assistance in facilitating interactions with the Angolan National Public Health authorities.

The following reagent was obtained through BEI Resources, NIAID, NIH: Diagnostic Plasmid Containing the Small Subunit Ribosomal RNA Gene (18S) from *Plasmodium falciparum*, MRA-177, contributed by Peter A. Zimmerman. *Plasmodium ovale* positive controls were kindly provided by Bruno Pradines, UMR RITMES, Marseille, France.

## Availability of data and materials

The data and materials that support the findings of this study are available from the corresponding author upon request. Sequences have been submitted to NCBI GenBank database.

## Declaration

This study was conducted in accordance with the Edinburgh revision of the Helsinki Declaration and was approved by Dr Filomeno Fortes from the National Malaria Control Program of the Ministry of Health of Angola.

## Consent for publication

Not applicable.

## Supporting information captions

S1 Table: Species-specific PCR identification of *Anopheles gambiae s.l.* and *An. funestus s.l*.

S2 Table: Abundance and diversity of *Anopheles* collected in 8 villages of Balombo in December 2023.

S3 Table: Abundance and diversity of *Anopheles* species by trap type and season in Balombo, 2024.

